# Huntington’s disease phenocopy syndromes revisited: a clinical comparison and next-generation sequencing exploration

**DOI:** 10.1101/2022.09.23.509161

**Authors:** CAM Koriath, F Guntotoi, P Norseworthy, E Dolzhenko, MA Eberle, DJ Hensman Moss, M Flower, H Hummerich, A Rosser, SJ Tabrizi, S Mead, E Wild

## Abstract

When the genetic test for the Huntington’s disease (HD) *HTT* expansion first became available almost 30 years ago, only 1% of patients tested negative. Since then, the test has become more accessible and the HD phenotype has expanded. More patients are being tested overall, and more negative tests are being received. These patients are deemed “HD phenocopy syndromes” (HDPC). In this study we established a current estimate for the prevalence of these patients. We also surveyed HD clinician experts on what would make them consider an HD test and compared both HD and HDPC patients to these expectations to decide whether they could be distinguished clinically; this proved impossible even when comparing symptom patterns. We re-analysed existing gene panel data for likely and potentially deleterious variants. Furthermore, we determined principles to prioritise patients for whole-genome sequencing (WGS). It was used to probe a 50 patient strong subcohort of HD phenocopy syndromes for known causes of HD-like and other neurodegenerative disease, identifying one *ATXN1* expansion using ExpansionHunter^®^. This was a small genetic substudy and therefore unsurprisingly no other known deleterious variants could be identified as in these cryptic understudied syndromes. Novel variants in known genes and variants in genes not yet linked to neurodegeneration may play an outsized role.

## Introduction

Huntington’s disease (HD), which is typically defined by a progressive triad of movement, cognitive, and psychiatric symptoms^1^, is the one of the commonest fatal, autosomal dominant, adult onset neurodegenerative disorders with a prevalence of at least 12.4 per 100,000 people^2,3^. Chorea is often considered the defining feature of HD, but motor symptoms can range from hyperkinetic to hypokinetic and often both. In addition, HD patients may experience a diverse range of symptoms, which may include cognitive impairment, anxiety, and depression even before the onset of unequivocal motor extrapyramidal symptoms^4^. Recent evidence suggests that the HD repertoire of both motor and non-motor manifestations may be caused by disruption of striatal gating through the direct and indirect pathways^5,6^. The heterogeneous presentation may render the clinical diagnosis difficult. Historically, when the test for the expansion in exon 1 of the Huntington’s gene (*HTT*)^7^ first became available only approximately 1% of patients with a clinical diagnosis of HD tested negative for the *HTT* expansion^8^. However, more recently the low cost and ready availability of the HD test is thought to have increased the negative test rate as clinicians may wish to exclude the disorder even if the syndrome is atypical^9^; a proportion of those in whom HD is suspected therefore do not carry the pathognomonic mutation. These patients are said to have an HD phenocopy (HDPC) syndrome and range from those mimicking “classical” HD exactly, to those with partially overlapping clinical features^10–12^. The differential diagnosis is wide even when only considering genetic causes^1^: expansion mutations in *C9orf72* have been identified as the most frequent identified genetic cause^10^ of HDPC syndromes, while spinocerebellar ataxia 17 (SCA17),, Huntington’s-like 2 (HDL2), and Friedreich’s ataxia (FRDA) have been found more rarely^12^. HDPC syndromes are an excellent example of unclear neurodegenerative disease presenting with symptoms spanning cognitive, psychiatric and motor disorders, promising insights into the genetic causes linking common functional categories or biological pathways - as has been previously studied in related neurodegenerative diseases like Alzheimer’s disease (AD) or frontotemporal dementia (FTD)^5^. Based on increased knowledge of what (variable) symptoms HD may present with^3^ and the hypothesis of increased HD testing, we surmise that HDPC syndromes are now more common than when the HD test first became available but that the original heuristic definition of “patients referred for HD testing by a clinician experienced in HD and who tests negative for the HD gene mutation” continues to be valid. We undertook a survey of expert opinion and practice and compared them to the clinical picture of patients referred for HD testing at the Laboratory of Neurogenetics at Queen Square, London, UK. Furthermore, we employed next-generation sequencing to explore the prevalence of deleterious mutations in known dementia genes and known HDPC causes in the UCL HDPC cohort.

## Methods

In order to compare the phenotype of HD and HDPC patients and how they match up with clinical expectations, 130 experienced neurologists and neurogeneticists from the European Huntington’s disease network (EHDN) were invited to take part in a 10-question online survey^13^ via email; email addresses of clinical leads were identified from the EHDN website, at least one clinician, all where possible, was contacted for each site, which were located in Europe and the USA. The 52 answer sets were anonymised and analysed using Microsoft Office Excel© and SPSS26.

In addition, clinical records of 151 patients from Neurogenetics clinics at the National Hospital for Neurology and Neurosurgery (NHNN) and Great Ormond Street Hospital between 2016 and 2018 were examined with regards to patients’ clinical symptoms at the time of genetic testing. In the case of pre-symptomatic HD testing of patients with a known positive genetic HD test result in the family, symptoms reported at the appointment following the onset of motor symptoms were recorded instead (see Figure 1). Results were analysed using SPSS26 (Fisher’s exact test) and R (logistical regression, conditional probabilities).

**Figure 1:**
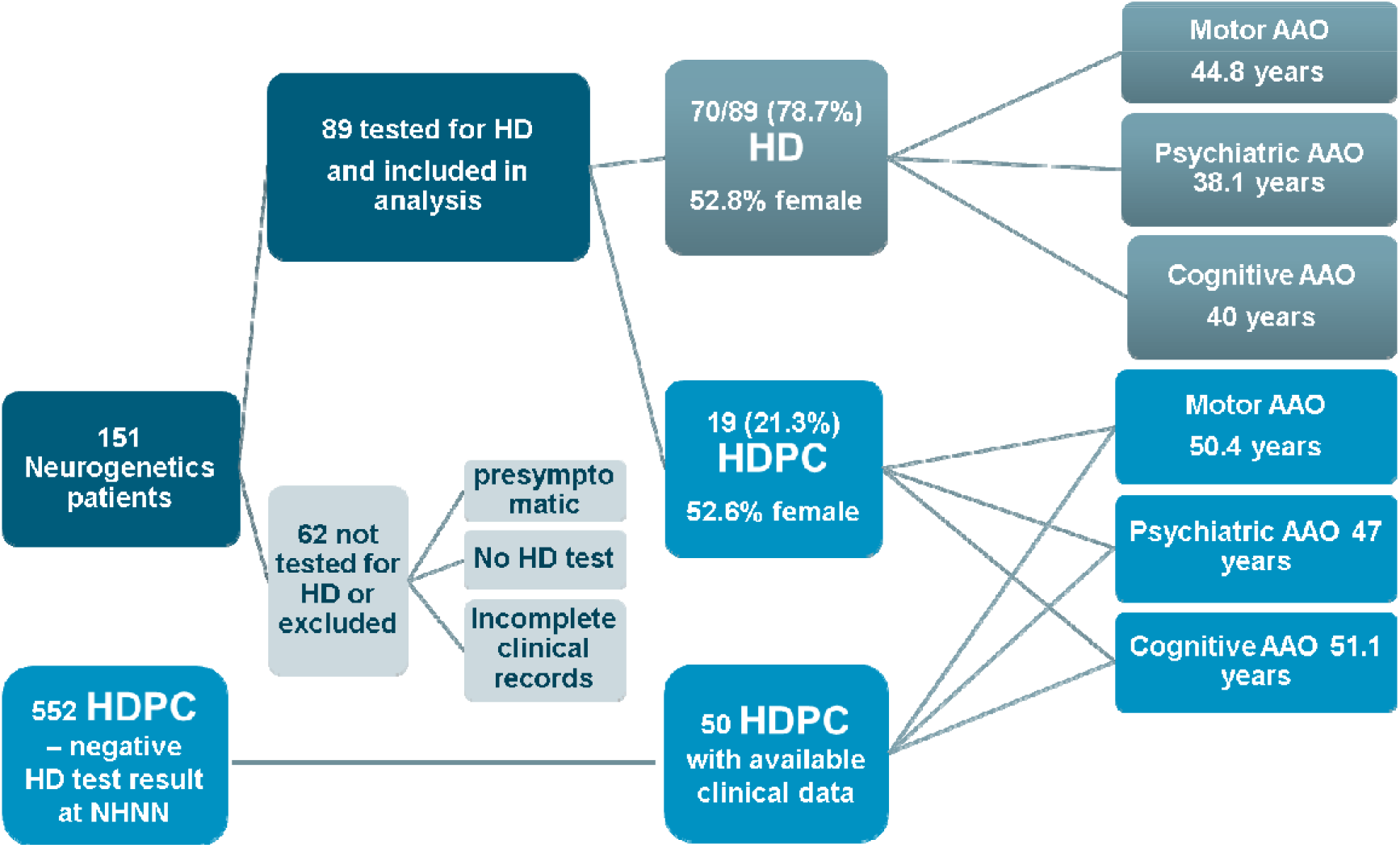
Patients from two clinics for Neurogenetics counselling and HD testing. 151 patients from two clinics for Neurogenetics were included in the analysis. 89 patients had an HD test; in 21.3% of cases tested for HD, the test result was negative. Ages at onset of motor symptoms were similar in patients with both a positive and a negative HD test result, but ages at onset of psychiatric and cognitive symptoms varied more. This analysis did not discriminate on the basis of a known family history of HD, i.e. whether testing was carried out pre-symptomatically. To improve statistical power, 50 additional Huntington’s disease phenocopy (HDPC) patients from the UCL HDPC genetic cohort for whom clinical data was available were added to the analysis.

For better statistical comparability and applicability, the Neurogenetics clinic HDPC cohort was supplemented with 50 of the most recently included patients from the UCL genetic cohort who were seen in other Neurology clinics at NHNN as well as other London hospitals and underwent genetic testing at NHNN. The additional genetic cohort cases were selected based on the availability of sufficient clinical information and whose HD test dates fall within the date range 2011 to 2015 (when the majority of the cases seen in the Neurogenetics Clinic were tested). Electronic patient records were reviewed for clinical information; cases with incomplete patient records or patients included in the Neurogenetics clinic cohort were excluded. We then undertook a deeper analysis of our previously-studied dementia gene panel, which was applied to 2784 patient and 457 control samples, including 552 HDPC patients, to examine the variants detected in this cohort^14^.

Furthermore, 50 HDPC cases were selected for whole genome sequencing (WGS) from the UCL HDPC cohort^10^ based on: how HD-like they were (HDPC score, adding one point for the presence of a movement disorder, cognitive decline, and psychiatric problems, respectively), the strength of their family history (FHx, encoded as the Goldman score)^15,16^, their age at onset, whether neuropathological data was available, and the number of years since they were last seen in clinic (see Table 1)^17^.

**Table 1:** Ranking and weighting factors for the selection of HD phenocopy samples for whole-genome sequencing. Each category of the criteria for each sample is awarded a number of points; these are then multiplied by the respective criterion’s weighting factor and added up to generate a total score for each sample. The maximum achievable score in this ranking is 100 points.

| Criteria | Category | Points awarded |
| --- | --- | --- |
| HD phenocopy score | 3 | 30 |
|  | 2 | 20 |
|  | 1 | 5 |
| Goldman score | 1 | 20 |
|  | 2 | 10 |
|  | 3 | 4 |
|  | 3.5 | 2 |
|  | 4 | 0 |
| Age at onset | ≤35 | 30 |
|  | ≤45 | 20 |
|  | ≤55 | 10 |
|  | ≤65 | 5 |
| Neuropathology available? | Yes | 15 |
|  | No | 0 |
| Years since seen in clinic | Last 2 years | 5 |
|  | Last 3 years | 2 |
|  | Last 5 years | 1 |
| Maximum Total |  | 100 |

The cohort has been previously screened for deleterious variants in genes known to be associated with dementia, APP, *CHMP2B, CSF1R, DNMT1, FUS, GRN, HTRA1, ITM2B, MAPT, NOTCH3, PRNP, PSEN1, PSEN2, TARDBP, TREM2, TYROBP* and *VCP*, as well as *C9orf72* expansions and *PRNP* octapeptide repeats^14^. No genetic or neuropathological diagnoses were available for the cases that were ultimately selected for WGS. Samples were sequenced on a commercial basis at Edinburgh Genomics: Clinical Genomics using Illumina SeqLab and Genologics Clarity LIMS X Edition (read length 150bp, average coverage 39.7x) and aligned to the Genome Reference Consortium Human Build 38 (GrCh38). Many known HD phenocopy syndromes are other expansion disorders; all samples were therefore screened using ExpansionHunter^®18^ for known expansion- and duplication-related causes of neurodegenerative disease in *AR, ATN1, ATXN1, ATXN2, ATXN3, ATXN7, ATXN10, C90RF72, CACNA1A, CBL, CSTB, DMPK, DMPK, FMR1, FXN, HTT, JPH3* and *PPP2R2B*. Samples without detected expansions were analysed using Ingenuity^®^ Variant Analysis™ software (IVA™, www.qiagenbioinformatics.com, version 5.4.20190308, Ingenuity Systems). Variants found at a minimum frequency of 0.1% in healthy public genomes such as Gnomad^19^ were excluded from the analysis; only variants with high confidence in the variant call (based on call quality, read depth, allele fraction, and location outside the top 5% of most exonically variable 100base windows and top 1% most exonically variable genes in healthy public genomes) and predicted to be deleterious, likely deleterious, or of uncertain significance were included, as well as those predicted to cause either gain-of-function (GoF) or loss-of-function (LoF) of a gene. A number of subsequent filters were explored including variants linked to neurological disease, variants predicted to be deleterious by the IVA^®^ algorithm, and variants linked to cell functioning in HD as per the IVA^®^ algorithm.

Informed consent for genetic studies was obtained from all participants. Ethical approval to undertake these analyses was given by the local NHNN/ION ethics committee.

## Results

### Survey

52 HD specialists responded to the survey; most had been working with HD and HDPC patients for a long time (mean 18.4 years, range 3-36 years) and saw many patients per month (mean 17 patients, range 1-50). They all considered chorea, and to a lesser extent cognitive slowing, executive dysfunction, irritability, gait abnormalities/falls, and dystonia indicative of HD, but neuropathy, limb weakness, ataxia, pain, tremor, and hallucinations suggestive of a different syndrome (Table S-1, Q5 and Q7). However, a substantial proportion of HD experts (10/52, 19.23%) felt that no ancillary symptom would make them expect a negative HD test result in the presence of a symptom calling for an HD test (Table S-1, Q10). Still, in the absence of a positive family history, only chorea, with or without additional symptoms, was deemed sufficiently suggestive to warrant testing for most (Table S-1, Q8). The symptoms most expected in HD included chorea, irritability, a dysexecutive syndrome, apathy, falls, gait abnormality, cognitive slowing, dystonia, and depression. The symptoms least expected in HD were neuropathy, limb weakness, pain, and tremor. (Q4 and Q6, Chi-Square tests, Bonferroni-corrected for multiple comparisons, see Table 2).

**Table 2:**
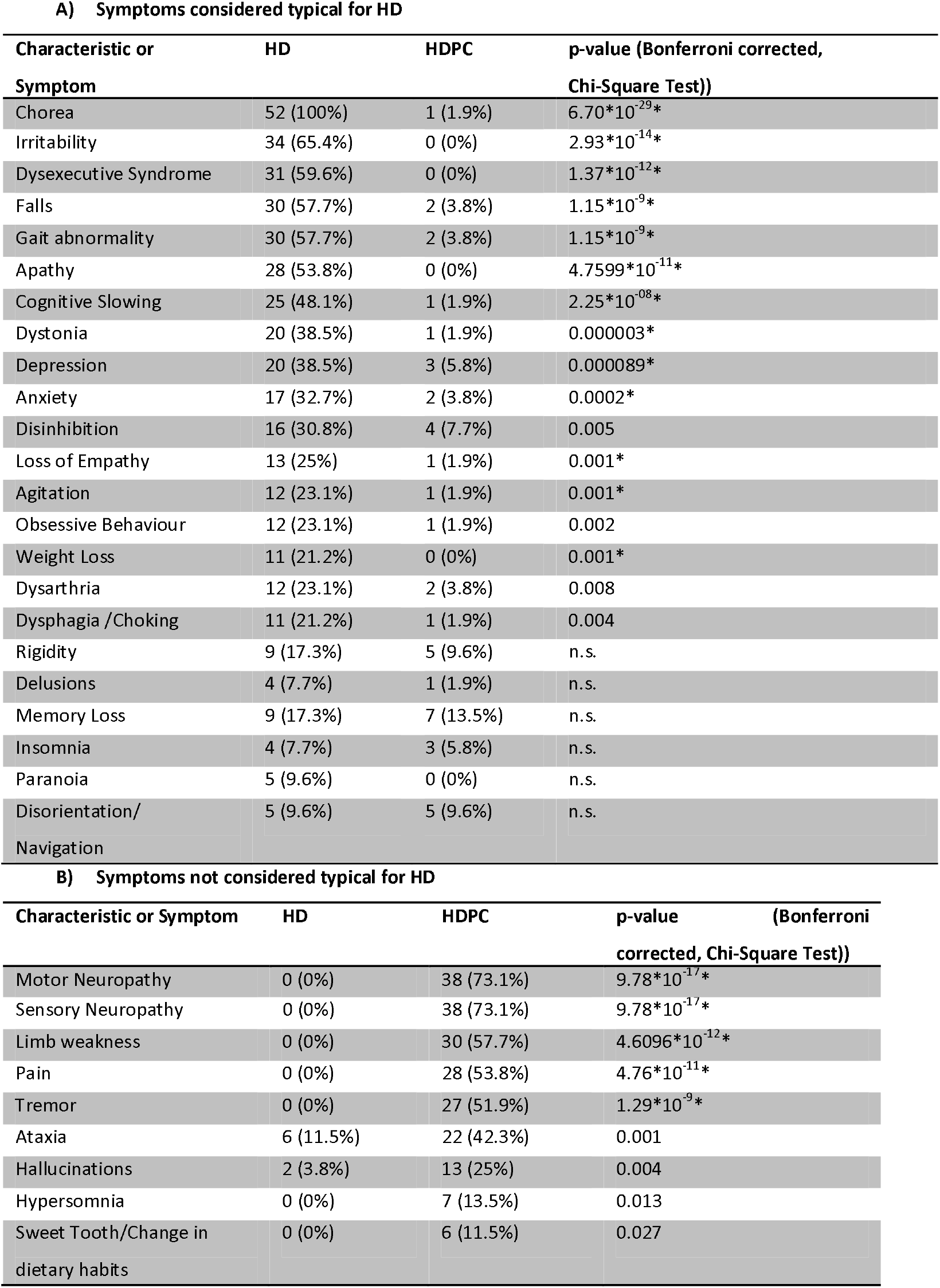
Symptoms expected to be typical for HD or not. P-values marked * remain significant at 0.05 after correcting for multiple testing using the Bonferroni method

### Clinical presentation at testing

151 patients seen in Neurogenetics clinics were analysed. Of 89 clinic patients tested for HD, 70 (78.7%) were found to carry the *HTT* expansion, while 19 patients (21.3%) did not (Figure 1). 62 patients were excluded for not having had an HD test, incomplete records, or no symptoms during follow-up of a presymptomatic test. Assuming comparable rates between centres, and given the estimated prevalence of HD of 10.6-13.7/100,000^20^, the prevalence of HDPC syndromes, i.e. patients with a movement disorder with cognitive and neuropsychiatric symptoms is calculated at 2.3-2.9/ 100,000 population. For three HDPC patients a diagnosis was confirmed during follow-up: one Parkinson’s disease with a typical dopamine transporter PET scan and two genetically confirmed spino-cerebellar ataxias (SCA 1 and SCA 17). 52.8% of HD and 52.6% of HDPC patients were female.

In order to improve statistical power, in addition to the 89 patients from Neurogenetics clinic who underwent genetic testing, an additional 50 HDPC cases were included in the comparison bringing the total to 139 cases – 70 HD and 69 HDPC cases (Figure 1 and Table 3). On average, HD patients were slightly younger at motor onset (mAAO) than HDPC patients; motor onset defines the onset of HD, however, many HD patients experienced much earlier psychiatric and cognitive symptoms, HPC patients less so. Initial presentation in clinic was very similar for HD and HDPC patients with symptoms including chorea, depression, and memory loss. Tremor, dystonia, and disinhibition were more common in HDPC, while dysphagia/choking, dysarthria, and insomnia were more frequently seen in HD. Weight loss was less specific than could have been expected based on survey results. While some oculomotor problems, tongue chorea, and motor impersistence can be very sensitive for HD, they were not well documented; no United HD Rating Scale (UHDRS) score was recorded. After Bonferroni’s correction for multiple testing, only family history was significantly different between the cohorts (Chi-Square-test, p=1.5058*10^-11^). A logistical regression test confirmed the differentiating effect of the Goldman score (p= 0.000009) between the HD and HDPC cohorts, but the individual GS categories did not reach significance. Having a strong autosomal (GS1) and having no relevant family history (GS4) was therefore almost reversed between the two cohorts; most HD patients had a strong known family history of HD, often with previously positive tests in the family, while most HDPC patients had a no established diagnosis in any known affected relatives.

**Table 3:** Characteristics of the HD/HDPC cohort. Symptom percentages are based on their being listed in the clinical notes and letters. Where possible a separate AAO for motor, cognitive and psychiatric symptoms was established from the notes (AAO Mot, AAO Cog, AAO Psy, respect.). In addition, the HDPC score was calculated for each patient based on whether they were displaying any symptoms in the cognitive, psychiatric and motor domains (cognitive, psychiatric, and motor HDPC score, respectively); this was used as a measure of how HD-like patients’ clinical presentations were in terms of affecting different functional domains.

| **A) Characteristics of the cohort** |  |  |  |
| --- | --- | --- | --- |
| **Characteristic or Symptom** | **HD (%HD)** | **HDPC (%HDPC)** | **p-value (uncorrected Chi-square/t-test)** |
| **Numbers** | 70 | 69 |  |
| **Male** | 33 (47.1%) | 29 (42.0%) | n.s. |
| **GS** | 2 (median) | 4 (median) | n.s. |
| **AAO Mot** | 44.9 years | 50.4 years | n.s. |
| **AAO Cog** | 40.0 years | 51.1 years | n.s. |
| **AAO Psy** | 37.4 years | 47.0 years | n.s. |
| **Total HDPC + FH score** | 2.9 | 2.6 | n.s. |
| **Psychiatric HDPC score** | 0.7 | 0.6 | n.s. |
| **Motor HDPC score** | 0.9 | 0.9 | n.s. |
| **Total HDPC score (noFHx)** | 2.1 | 2.2 | n.s. |
| **B) Symptoms suggestive of HD** |  |  |  |
| **Characteristic or Symptom** | **HD (%HD)** | **HDPC (%HDPC)** | **p-value (uncorrected Chi-square/t-test)** |
| **Insomnia** | 24 (34.3%) | 8 (11.6%) | 0.002 |
| **Dysphagia /Choking** | 22 (31.4%) | 8 (11.6%) | 0.007 |
| **Dysarthria** | 24 (34.3%) | 12 (17.4%) | 0.033 |
| **Falls** | 23 (32.9%) | 14 (20.3%) | n.s. |
| **Irritability** | 23 (32.9%) | 14 (20.3%) | n.s. |
| **Depression** | 25 (35.7%) | 19 (27.5%) | n.s. |
| **Ocular Saccades** | 22 (31.4%) | 16 (23.2%) | n.s. |
| **Agitation** | 6 (8.6%) | 3 (4.3%) | n.s. |
| **Apathy** | 7 (10%) | 4 (5.8%) | n.s. |
| **Weight Loss** | 8 (11.4%) | 5 (7.2%) | n.s. |
| **Rigidity** | 11 (15.7%) | 8 (11.6%) | n.s. |
| **Anxiety** | 18 (25.7%) | 15 (21.7%) | n.s. |
| **Ocular Smooth Pursuit** | 17 (24.3%) | 14 (20.3%) | n.s. |
| **Gait abnormality** | 38 (54.3%) | 35 (50.7%) | n.s. |
| **Chorea** | 48 (68.6%) | 45 (65.2%) | n.s. |
| **Hypersomnia** | 1 (1.4%) | 0 (0%) | n.s. |
| **Bradykinesia** | 12 (17.1%) | 11 (15.9%) | n.s. |

C) Symptoms less suggestive of HD
| Characteristic or Symptom | HD (%HD) | HDPC (%HDPC) | p-value (uncorrected Chi-square/t-test) |
| --- | --- | --- | --- |
| Dystonia | 7 (10%) | 23 (33.3%) | 0.001 |
| Tremor | 3 (4.3%) | 18 (26.1%) | 0.00031 |
| Disinhibition | 0 (0%) | 15 (21.7%) | 0.000012 |
| Cognitive HDPC score | 0.5 | 0.6 | n.s. |
| Cognitive Slowing | 17 (24.3%) | 22 (31.9%) | n.s. |
| Dysexecutive Syndrome | 22 (31.4%) | 26 (37.7%) | n.s. |
| Limb weakness | 0 (0%) | 4 (5.8%) | n.s. |
| Spasticity | 0 (0%) | 4 (5.8%) | n.s. |
| Supranuclear Palsy | 1 (1.4%) | 5 (7.2%) | n.s. |
| Myoclonus | 3 (4.3%) | 6 (8.7%) | n.s. |
| OcularRangeofMovement | 3 (4.3%) | 6 (8.7%) | n.s. |
| Paranoia | 1 (1.4%) | 4 (5.8%) | n.s. |
| Loss of Empathy | 0 (0%) | 3 (4.3%) | n.s. |
| Sensory Neuropathy | 0 (0%) | 3 (4.3%) | n.s. |
| Pain | 0 (0%) | 3 (4.3%) | n.s. |
| Sweet Tooth/Change in dietary habits | 0 (0%) | 3 (4.3%) | n.s. |
| Obsessive Behaviour | 3 (4.3%) | 5 (7.2%) | n.s. |
| Disorientation/Navigation | 2 (2.9%) | 4 (5.8%) | n.s. |
| Hallucinations | 1 (1.4%) | 3 (4.3%) | n.s. |
| Ataxia | 12 (17.1%) | 13 (18.8%) | n.s. |
| Asymmetry | 1 (1.4%) | 2 (2.9%) | n.s. |
| Babinski | 1 (1.4%) | 2 (2.9%) | n.s. |
| Delusions | 2 (2.9%) | 3 (4.3%) | n.s. |
| Bladder/ Bowel Dysfunction | 0 (0%) | 1 (1.4%) | n.s. |
| Cortical Blindness | 0 (0%) | 1 (1.4%) | n.s. |
| Nystagmus | 0 (0%) | 1 (1.4%) | n.s. |
| Memory Loss | 27 (38.6%) | 27 (39.1%) | 1 |
| Lost Reflexes | 0 (0%) | 0 (0%) | N/A - constant |
| Motor Neuropathy | 0 (0%) | 0 (0%) | N/A - constant |

**Table 4:**
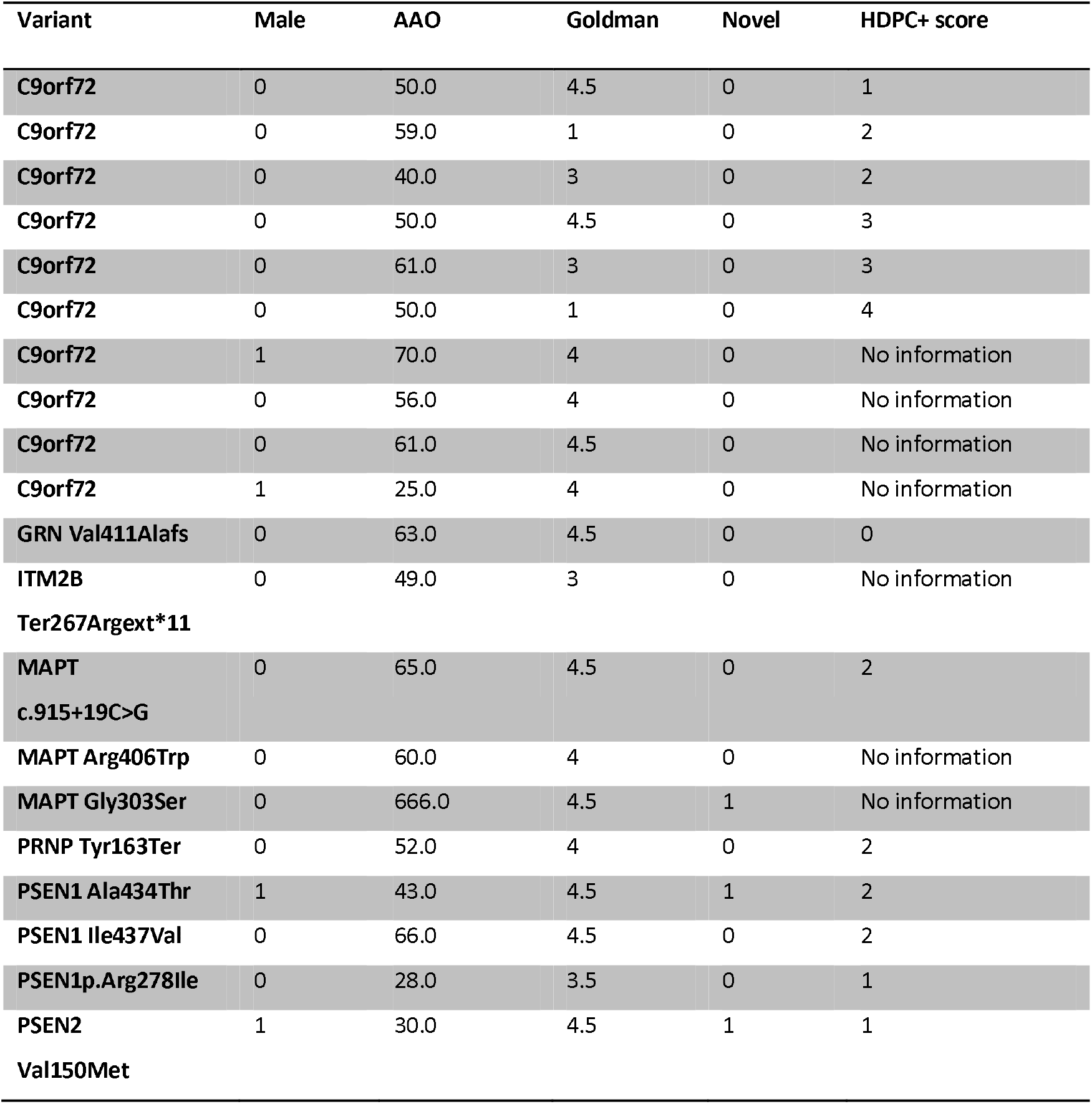
Summary of at least likely deleterious variants identified in HDPC patients in the gene panel dataset. Shown are the variants and details of patients in whom they were identified, including sex, age at onset, the Goldman score as a measure of family history, as well as the HDPC+ score (HDPC score plus positive family history) as a measure of how HD-like these patients were. Missing data is displayed 666.

Given the variability in presentation, however, comparing symptom patterns by calculating the conditional probability of two given symptoms co-occurring may offer much insight into the eventual diagnosis and future disease course (Figure 2). While both HD and HDPC syndromes present with subcortical dementia, cognitive slowing and executive dysfunction, combined dysarthria and memory loss early on in the disease course in HD (conditional probability 0.71/0.63) highlight its links between motor and cognitive symptoms; early insomnia and memory loss also appear suggestive of HD. Furthermore, HD patients combine depression and irritability (conditional probability 0.56/0.61) more often than HDPC patients. HDPC patients typically show strong connections between related symptoms, such as disinhibition and irritability (conditional probability 0.64/0.6), as well as executive dysfunction and cognitive or memory loss (conditional probability 0.77/0.65 and 0.69/0.67, respectively). However, rates of chorea were similar in both cohorts.

**Figure 2.**
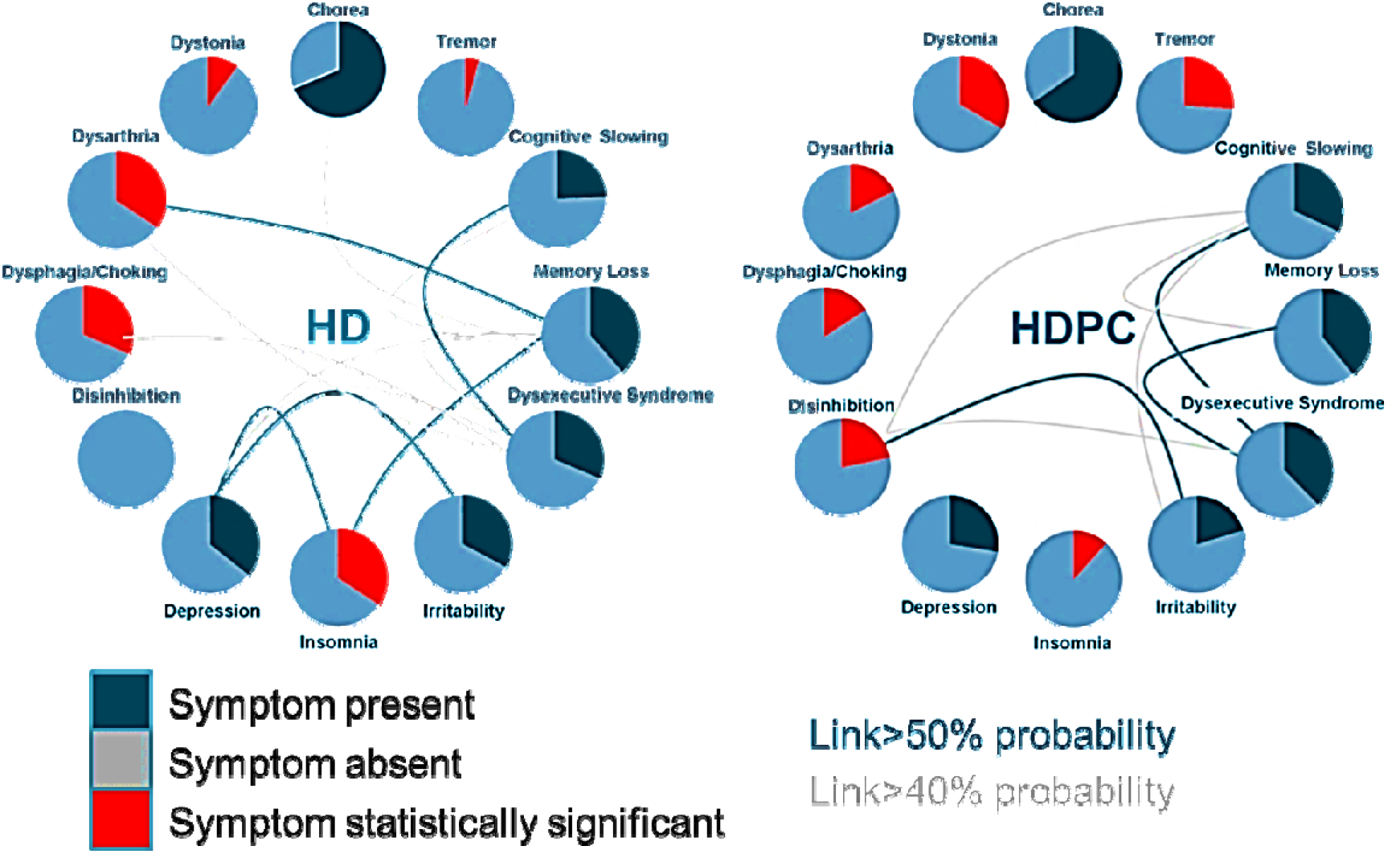
HD and HDPC display different patterns of concurrent symptoms: Each symptom circle shows the percentage of patients in each group presenting with a given symptom; the circle is highlighted in red if the symptom differed significantly between the two groups (p≤0.05, logistical regression). Given the high number of included symptoms, none of the results withstood Bonferroni’s correction for multiple comparisons, but they results are interesting from an exploratory vantage point. The likelihood of a patient presenting with any given symptom pair was calculated using conditional probabilities, which express the likelihood, that one symptom occurs given the presence of another in any given patient. If the likelihood is bilaterally equal or higher than 40%, a faint grey line connects the two symptoms, if it is equal or higher than 50%, the connecting line is emphasized in dark grey. All statistical analyses were done in R version 3.5.1 (https://www.rproject.org/) using the base package and custom written scripts.

### Genetic sequencing of HDPC patients

In the gene panel dataset, 319/552 (57.8%) HDPC patients were female, with an average AAO of any symptoms of 55.2 years. 22 patients (4%) had a GS of 1, 9 patients (1.6%) had a GS of 2, 16 patients (2.9%) had a GS of 3, 74 patients (13.4%) had a GS of 3.5, 207 patients (37.5%) had a GS of 4, and 224 patients (40.6%) had a GS of 4.5. At least some clinical information was available for 361 patients, who had a mean HDPC+ score (plus positive family history) of 1.4/4. 77.1% of these patients had a documented movement disorder, 37.4% had cognitive decline and in 12.3% of patients psychiatric symptoms had been noted.

Our reanalysis of the gene panel dataset^14^ focused on these HDPC cases revealed deleterious and likely deleterious variants in 20 (3.62%) HDPC cases compared to 3 (0.66%) in controls (p= 0. 0016, Chi-Square test). Classified based on previously published criteria^14^, deleterious and likely deleterious variants found in HDPC cases included *C9orf72* expansions in 10 patients, *MAPT* Arg406Trp, *MAPT* Gly303Ser, *MAPT* c.915+19C>G, *GRN* Val411Alafs, *ITM2B* Ter267Argext*11, *PRNP* Tyr163Ter, *PSEN1* Ala434Thr, *PSEN1* Arg278Ile, *PSEN1* Ile437Val, and *PSEN2* Val150Met. Three mutations were novel and have not been described before. No concurrent deleterious variants were identified in this cohort. Average AAO of the 20 patients with deleterious or likely deleterious mutations was 82.2 years old, mean Goldman score was 3.75, 17 patients were female. 7 patients had a documented movement disorder, 6 were known to have cognitive symptoms and 3 had psychiatric symptoms. No clinical information was available for 7 patients.

In addition to the deleterious and likely deleterious variants, 22 (4.0%) variants classified as potentially deleterious were observed in HDPC cases, compared to 6 (1.31%) in controls (p= 0.01730, Chi-Square test). Whilst the available data is not sufficient to classify variants in the potentially deleterious category as disease-causing, they were found in excess in cases vs. controls in the whole dataset (p=0.0039, Odds Ratio: 3.2, 95% CI (1.39, 7.28)) and most likely to be identified in HDPC patients (p=0.00012; Odds Ratio: 4.63, 95% CI (1.92, 11.16)) compared to AD (p=0.0129; Odds Ratio: 2.88, 95% CI (1.21, 6.89)) or FTD patients (p=0.0099; Odds Ratio: 3.06, 95% CI (1.25, 7.47)). In contrast, novel variants reaching the threshold for deleterious or likely deleterious variants were not observed significantly more frequently in the HDPC cohort compared to controls (logistical regression, p>0.05). Potentially deleterious variants observed in the HDPC dataset included *CSF1R* Arg549Cys, *FUS* Gly227_Gly229del, *FUS* Gly229Ser, *FUS* Pro431Leu, *GRN* Arg478His, *GRN* Asp33Glu, *GRN* Gly148Arg, *GRN* Pro166Leu, *NOTCH3* Asp1869Gly, *PSEN1* His46Tyr, *PSEN1* Ile148Val,, *PSEN2* Asp431Glu, *SERPINI1* Leu307Ser, *SERPINI1* Ser142Gly, *SERPINI1* Val288Ile, *SQSTM1* Thr339Ile, *SQSTM1* Val240Ala, *SQSTM1* Val271Ile, as well as *VCP* c.1359+8C>T, *VCP* c.-221_-216delGCTGCC, *VCP* c.-254_-253insCGCTGCCGCTGCCGCTGC.

Furthermore, in order to explore other potential genetic causes in an unbiased manner, a subset of 50 patients were selected for WGS from the genetic UCL cohort. Of these, 29 patients (58%) were female with an average AAO of 51.2 years, a median GS of 3.5, and a mean HDPC+ score (plus positive family history) of 2.86/4. The average coverage per sample was 39.7x. 92% of patients presented with a movement disorder (56% with chorea), 76% had cognitive decline and 54% had psychiatric symptoms including behavioural and personality change, affective changes, as well as obsessive or psychotic symptoms (see Table S-2). Samples were screened for known trinucleotide repeat expansions using ExpansionHunter^®18^, which correctly identified a, since clinically confirmed, Ataxin 1 expansion (see Figure 3) known to cause SCA1. SCA1 is an established HD phenocopy^11,21^: the patient’s symptoms started at the age of 40 with gait and coordination problems; he had a positive family history, consistent with a diagnosis of SCA1. No other expansions in the genes known to be linked to expansion disorders were identified in any other samples.

**Figure 3::**
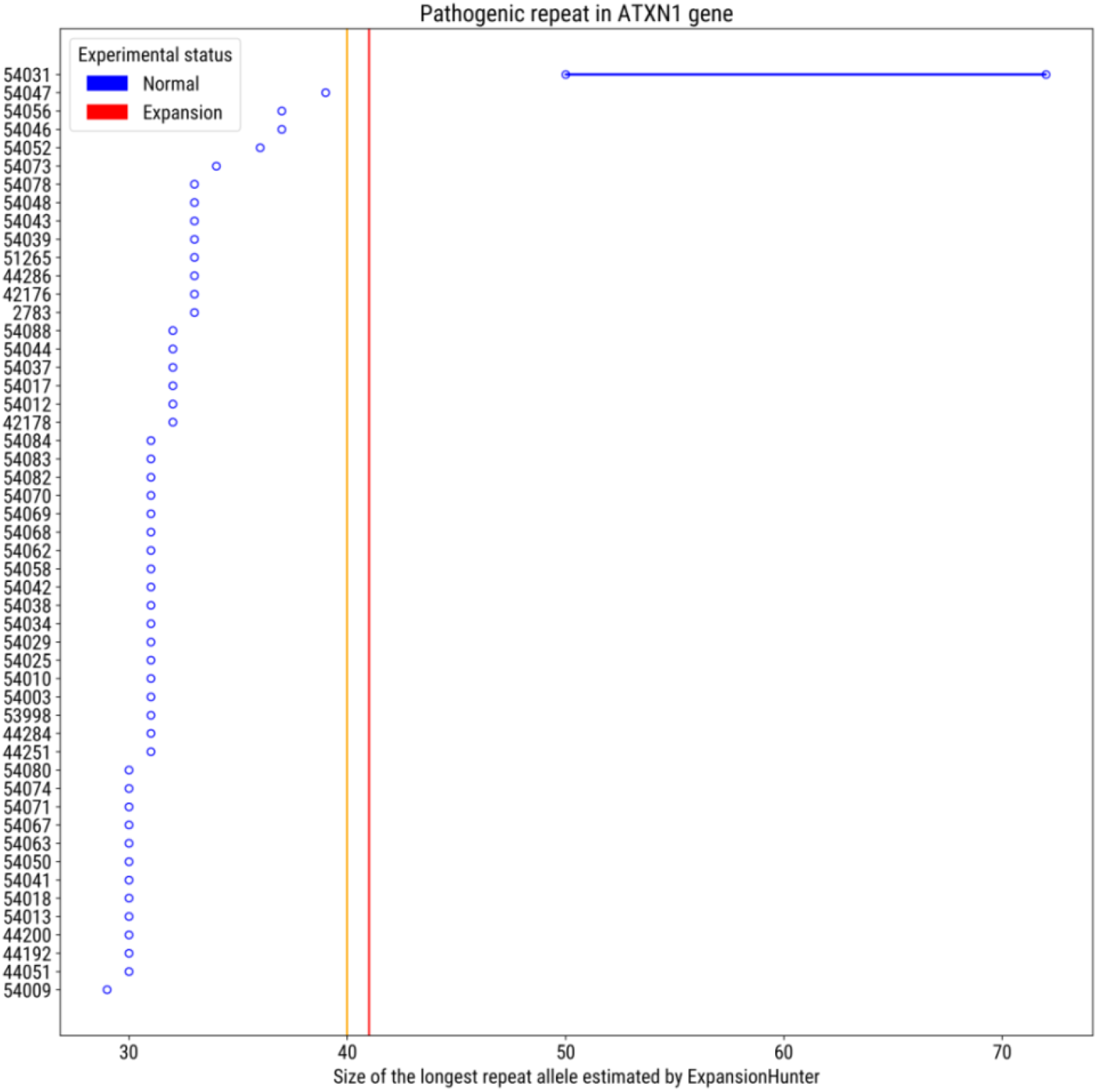
ExpansionHunter^®^ identifies a repeat expansion in the *ATXN1* gene. ExpansionHunter^®^ correctly identified an expansion in the *ATXN1* gene, as can be observed on this output from the programme. All but one sample were estimated to have repeat sizes well below the 39 CAG repeats found in patients with spino-cerebellar ataxia type 1 (SCA1).

Analysis of the 49 remaining samples in IVA™ showed an average of 5,334,908 variants per sample (range 5,230,677 to 6,331,549). Given the estimated penetrance of a causal variant observed once in this dataset and once on *GnomAD*^14,19,22^, only known deleterious and novel variants were considered for further analysis, but no more known deleterious variants could be identified.

## Discussion

Over the last 30 years, neurodegenerative syndromes have progressively been recognized as heterogeneous and multifactorial disorders as the number of known Mendelian causes of disease has increased. One of earliest clinically available tests was the *HTT* expansion test^7^, which was initially positive in 99% of clinically-diagnosed HD patients^8,23^. However, since then HD testing has become much more accessible, the breadth of the recognized HD phenotype has expanded, and it has become customary to test for HD even if a patient’s presentation may not be a typical case, leading to suggestions that some HDPC patients should not have been tested for HD at all because their symptoms were not sufficiently typical of HD^9^. We therefore sought to establish whether HD and HDPC syndromes can be distinguished clinically based on what clinical HD experts considered typical for HD, translating to when they may be inclined to order a genetic test, and what HD and HDPC patients’ symptoms in clinic were at the point of testing. Furthermore, we developed a system for prioritizing patients for WGS and analysed a subcohort of 50 patients for known deleterious mutations. While the aetiology of HD is well-established, it remains unclear how it translates to the diverse symptoms; a better understanding of what other genes cause a similar phenotype may aid in this process.

Based on the survey results, chorea was considered to be typical of HD, as were dystonia, executive dysfunction and cognitive slowing, apathy and depression. However, while the typical manifestations of HD are hyperkinetic and HD is the cause of 90% of genetic chorea^24^, it is a heterogeneous disease potentially presenting with relatively pure dystonic, ataxic, and psychiatric symptoms; as the disease progresses, gait and postural disturbance, as well as falls eventually follow^21^. In addition, most patients with juvenile onset, and some adult-onset patients, present with the hypokinetic *Westphal* variant, sometimes in combination with seizures, further broadening the disease spectrum^25^.

Our results show that HD and HDPC were generally impossible to distinguish in clinic based on single symptoms; noted differences in dystonia, dysarthria and dysphagia were minor and in some instances could also reflect differences in timing of presentation or documentation. Contrary to expert opinion, HD and HDPC patients presented with comparable levels of chorea, as well as executive dysfunction, apathy, gait abnormality and others.

Dystonia, which is considered a possible indicator of HD, was present in 33.3% of HDPC but only 10% of HD patients (not significant after correction for multiple testing). Dystonia is a typical feature of HD^26^, often in combination with chorea but sometimes the only one for some time^27^, and has been shown to detrimentally affect day-to-day functioning in HD patients more than chorea^28^, but pronounced dystonia is also common in HDL2, HDL3 and SCA17/HDL4, as well as in HDPC syndromes associated with *C9orf72*, neuroferritinopathy and brain calcifications^21^.

As predicted by the survey, insomnia, dysarthria, and dysphagia were also significantly more common in HD than in HDPC. Circadian disruption is a feature of many neurodegenerative diseases possibly affecting progression^29^; insomnia can have secondary effects on cognitive and emotional symptoms in HD^30^. In contrast, speech dysfunction has been demonstrated even in premanifest HD patients^31^ but little longitudinal data is available, while dysphagia can be used to track disease progression in HD^32^.

According to HD patients themselves, dysexecutive and cognitive symptoms have the greatest negative impact on their lives^33,34^ as consequently impaired decision-making also disrupts their social functioning long before their motor symptoms become debilitating^35^. However, recent evidence in FTD indicates that profiles of cognitive decline may subtly diverge depending on the genetic cause^36^.

Even before motor onset neuropsychiatric symptoms are very common in HD patients and increase in prevalence as the disease develops^37^. Apathy is highly prevalent in HD^38^ and was even found to predict the rate of cognitive decline in presymptomatic and manifest HD patients^39,40^. Depression, meanwhile, is very common in diverse neurodegenerative disease and responds poorly to current antidepressant treatment^41^.

Notably, almost 20% of survey respondents could not be influenced in their inner prediction of a patient’s test result even by very disparate symptoms, such as neuropathy, limb weakness, pain, tremor, and ataxia, when the need to test for HD based on a different concurrent symptom was perceived. However, in this study, neuropathy, limb weakness, and pain were indeed exclusively observed in HDPC patients, while tremor was seen significantly more often in HDPC than in HD patients. Neuropathy and ataxia may be associated with the spinocerebellar ataxias, Friedreich’s ataxia or mutations in the *POLG* or *TWNK* genes^10,11,42,43^, while limb weakness may be indicative of vascular disease^44,45^. Tremor has been described in *C9orf72*-linked HDPC syndromes^21^, and can also feature in SCA6 and SCA12^46^. Patients with neurodegenerative ataxias have much higher prevalences of tremor than HD patients, but in both cohorts the predominant type is essential tremor^9^. On closer examination, however, patients with a diagnosis of essential tremor often have undiagnosed conditions, such as dystonia and Parkinson’s disease, and may require a more detailed work-up^47^.

The clinical presentation of HD and HDPC patients did differ significantly when symptom combinations were analysed together. Relatively early orolingual involvement with dysarthria, dysphagia or choking in patients with cognitive problems or hyperkinetic limb movements appears suggestive of HD. This may in part be due to the composition of our cohort, since a recent study showed that dystonia and dysarthria were more common in HDL2 than HD patients, although associations were not analysed^48^. Furthermore, different types of dysarthria in HD have been shown to vary significantly in terms of general motor dysfunction^49^. HD patients may also suffer from primary deficits of language^50^, further confounding the motor and cognitive tasks of speech and language. Consistent with reported links between sleep disturbance, cognitive decline and neuropsychiatric symptoms in HD^51^, the combination of early insomnia, memory loss, and depression was more common in HD than in HDPC patients. While levels of depression and irritability were similar, their link was stronger in HD than in HDPC. Irritability has previously been associated with an early increase in motor symptoms in HD, and although this was not the case for depression, the authors did not comment on the use of antidepressant medication^52^.

The level of neuropsychiatric symptoms appears to be greater in HD patients with a hypokinetic-dominant motor presentation, rather than the chorea-dominant or mixed-motor phenotype^53^. Overall, symptoms in HDPC patients seemed to commonly be less diverse and associations were stronger between related symptoms, such as disinhibition and irritability. Cognitive and psychiatric symptoms were mostly observed with a similar frequency in the two cohorts, but disinhibition (rather than depression, apathy or anxiety) could point the clinician towards an HDPC syndrome; this was especially true in combination with irritability, cognitive slowing and executive dysfunction. Interestingly, apathy, which has previously been reported to be a core feature of HD^37^ and shown to present similarly to a cognitive symptom^54^, was not associated with any other symptoms in this analysis; this may be due to difficulties of a retrospective study such as gaps in the clinical documentation, or because of underlying patient heterogeneity.

In this study, HD expert clinicians’ expectations therefore mirrored most of the heterogeneous results gathered from the patient analysis, suggesting that while testing has expanded, so has the accepted clinical HD phenotype (based in turn on experience of patients with positive test results). Twenty-five years after the discovery of the *HTT* expansion, one of the strongest indicators of an HDPC syndrome rather than HD appears to be the lack of an HD family history, or indeed any relevant family history of cognitive decline, psychiatric problems or motor symptoms. Given the results in this patient series and the estimated prevalence of HD at 10.6 to 13.7/100,000^20^, rates of HDPC syndromes have increased to approx. 20% of suspected HD cases; the prevalence of cryptic HDPC syndromes can therefore be estimated at approx. 2.5/100,000.

According to the genetic testing algorithm^10^, patients with an HD-like presentation should first be tested for the *HTT* expansions, followed by the *C9orf72* and *TBP* expansions (responsible for 1.95%^10^ and 1.1%^11^ of HDPC cases, respectively). Specific symptoms or details from the history may justify testing for other conditions, such as seizures for dentatorubral-pallidoluysian atrophy (DRPLA) or African heritage for HDL2^10^. However, once a patient with an HDPC syndrome has tested negative for these conditions, the diagnostic rate plummets^14,55–58^, our data suggests that novel variants and different genes (or non-genetic causes) may play an outsize role in HDPC syndromes or potentially act as modifiers in HD. A different approach is therefore needed in order to offer HDPC patients the benefits of a diagnosis. In the HDPC patients included in the present study, the *HTT* and *C9orf72* expansions, *PRNP* octapeptide repeats and deleterious variants in 17 dementia genes had already been studied^14^ before WGS sequencing. In this previous gene panel dataset, the established deleterious variants were least likely to be identified in HDPC patients compared to patients with Alzheimer’s disease, Frontotemporal dementia or Prion disease. However, the frequently novel, potentially deleterious variants were most commonly observed in HDPC patients, who present a more heterogeneous patient group. Less well characterized variants in dementia genes may therefore confer atypical features; the phenotypic spectrum of genes already known to be associated with neurodegeneration may be broader than has been understood so far and more hitherto unrecognized deleterious variants in these genes are yet to be discovered.

In addition, to eliminate other causal trinucleotide expansions in the WGS subcohort, samples were examined using the ExpansionHunter^®^ programme, which identified one expansion in *ATXN1* in a patient in whom the diagnosis of SCA1 was confirmed in a clinical laboratory. HD, the HD-like disorders, and other neurodegenerative diseases with dementia and motor symptoms are caused by trinucleotide repeats^11,21,46^ and should be excluded. Considering the limits of fragment analysis, southern blot^59–61^, and long-read sequencing^62^, bioinformatics analysis methods, such as ExpansionHunter^®^, are one of the most accessible forms of screening WGS data for expansion disorders while also searching for SNPs and smaller indels^18^ with high sensitivity and specifity^63^ and are not restricted to PCR-free WGS datasets. Subsequently, the HDPC samples were analysed via the IVA™ online platform; however, no further known deleterious variants could be identified. Given that this was a small study in a very heterogeneous study population, it is hardly surprising that no pertinent, known deleterious variants were found. Novel and understudied variants may play an outsize role in these less well-established syndromes^17^ and will require further study; genes not previously established to cause disease may also be involved and will require further study and replication.

In summary, HD and HDPC syndromes were clinically indistinguishable in this study on the basis of single symptoms, in contrast to expert HD clinician’s expectations. HDPC disorders are likely to encompass highly heterogeneous patients because HD itself is one of the most heterogeneous genetic neurological diseases, but patterns of symptoms, particularly the combination of early orolingual involvement with a hyperkinetic movement disorder may be suggestive of HD. Our data suggests an outsize role for novel variants in known dementia genes in these cases.

The study of a subcohort using WGS found one *ATXN1* expansion known to cause SCA1, but no further known deleterious variants could be identified.

## Acknowledgements

CK gratefully acknowledges the support of the Leonard Wolfson Foundation and the CHDI Foundation. S.J.T. received grant funding for her HD research from the Medical Research Council UK, the Wellcome Trust, the Rosetrees Trust, Takeda Pharmaceuticals, Cantervale Limited, the NIHR North Thames Local Clinical Research Network, the UK Dementia Research Institute, the Wolfson Foundation for Neurodegeneration and the CHDI Foundation. SM is supported by the Medical Research Council (UK), the NIHR Queen Square Dementia Biomedical Research Unit and the NIHR Biomedical Research Centre at University College Hospitals NHS Foundation Trust. EJW reports grants from Medical Research Council, CHDI Foundation, and F. Hoffmann-La Roche Ltd; personal fees from Hoffman La Roche Ltd, Triplet Therapeutics, PTC Therapeutics, Takeda, Teitur Trophics and Vico Therapeutics. All honoraria for these consultancies were paid through the offices of UCL Consultants Ltd., a wholly owned subsidiary of University College London. University College London Hospitals NHS Foundation Trust has received funds as compensation for conducting clinical trials for Ionis Pharmaceuticals, Pfizer and Teva Pharmaceuticals. DHM is funded through the University of London Chadburn Lectureship programme.

## References

1. Ghosh, R. & Tabrizi, S.J. Clinical Features of Huntington’s Disease. Adv Exp Med Biol 1049, 1–28 (2018).

2. Rawlins, M. Huntington’s disease out of the closet? Lancet 376, 1372–3 (2010).

3. Bates, G.P. et al. Huntington disease. Nat Rev Dis Primers 1, 15005 (2015).

4. Ghosh, R. & Tabrizi, S.J. Huntington disease. Handb Clin Neurol 147, 255–278 (2018).

5. Nair, A., Aziz, N.A., Rutledge, R., Rees, G. & Tabrizi, S.J. Relationship between Distinct Motor Symptoms and Apathy in Huntington’s Disease: Clues to Mechanism. Journal of Neurology Neurosurgery and Psychiatry 89, A56–A56 (2018).

6. Nair A, R.A., Gregory S, Rutledge RR, Rees G, Tabrizi SJ. Imbalanced basal ganglia connectivity is associated with motor deficits and apathy in Huntington’s disease. Brain. (2021).

7. A novel gene containing a trinucleotide repeat that is expanded and unstable on Huntington’s disease chromosomes. The Huntington’s Disease Collaborative Research Group. Cell 72, 971–83 (1993).

8. Andrew, S.E. et al. Huntington disease without CAG expansion: phenocopies or errors in assignment? Am J Hum Genet 54, 852–63 (1994).

9. Feigin, A. & Talbot, K. Expanding the genetics of huntingtonism. Neurology 82, 286–7 (2014).

10. Hensman Moss, DJ. et al. C9orf72 expansions are the most common genetic cause of Huntington disease phenocopies. Neurology 82, 292–9 (2014).

11. Wild, EJ. et al. Huntington’s disease phenocopies are clinically and genetically heterogeneous. Mov Disord 23, 716–20 (2008).

12. Wild, EJ. & Tabrizi, SJ. Huntington’s disease phenocopy syndromes. Curr Opin Neurol 20, 681–7 (2007).

13. Koriath, C., Wild, E., Tabrizi, S. Huntington’s Disease and HD-like syndromes. Vol. 12/04/2019.

14. Koriath, C. et al. Predictors for a dementia gene mutation based on gene-panel next-generation sequencing of a large dementia referral series. Mol Psychiatry (2018).

15. Goldman, J.S. et al. Comparison of family histories in FTLD subtypes and related tauopathies. Neurology 65, 1817–9 (2005).

16. Rohrer, J.D. et al. The heritability and genetics of frontotemporal lobar degeneration. Neurology 73, 1451–6 (2009).

17. Koriath, C. Univerity College London (2020).

18. Dolzhenko, E. et al. Detection of long repeat expansions from PCR-free whole-genome sequence data. Genome Res 27, 1895–1903 (2017).

19. Karczewski, K.J. et al. The mutational constraint spectrum quantified from variation in 141,456 humans. Nature 581, 434–443 (2020).

20. McColgan, P. & Tabrizi, S.J. Huntington’s disease: a clinical review. Eur J Neurol 25, 24–34 (2018).

21. Schneider, S.A. & Bird, T. Huntington’s Disease, Huntington’s Disease Look-Alikes, and Benign Hereditary Chorea: What’s New? Mov Disord Clin Pract 3, 342–354 (2016).

22. Minikel, E.V. et al. Quantifying prion disease penetrance using large population control cohorts. Sci Transl Med 8, 322ra9 (2016).

23. Kremer, B. et al. A worldwide study of the Huntington’s disease mutation. The sensitivity and specificity of measuring CAG repeats. N Engl J Med 330, 1401–6 (1994).

24. Schneider, S.A., Walker, R.H. & Bhatia, K.P. The Huntington’s disease-like syndromes: what to consider in patients with a negative Huntington’s disease gene test. Nat Clin Pract Neurol 3, 517–25 (2007).

25. Stout, J.C. Juvenile Huntington’s disease: left behind? Lancet Neurol 17, 932–933 (2018).

26. van de Zande, N.A. et al. Clinical characterization of dystonia in adult patients with Huntington’s disease. Eur J Neurol 24, 1140–1147 (2017).

27. Andriuta, D., Tir, M., Godefroy, O. & Krystkowiak, P. Huntington’s Disease Revealed by Familial Cervical Dystonia. Mov Disord Clin Pract 3, 415–416 (2016).

28. Carlozzi, N.E. et al. How different aspects of motor dysfunction influence day-to-day function in huntington’s disease. Mov Disord 34, 1910–1914 (2019).

29. Abbott, S.M. & Videnovic, A. Chronic sleep disturbance and neural injury: links to neurodegenerative disease. Nat Sci Sleep 8, 55–61 (2016).

30. Herzog-Krzywoszanska, R. & Krzywoszanski, L. Sleep Disorders in Huntington’s Disease. Front Psychiatry 10, 221 (2019).

31. Chan, J.C.S., Stout, J.C. & Vogel, A.P. Speech in prodromal and symptomatic Huntington’s disease as a model of measuring onset and progression in dominantly inherited neurodegenerative diseases. Neurosci Biobehav Rev 107, 450–460 (2019).

32. Manor, Y. et al. Dysphagia characteristics in Huntington’s disease patients: insights from the Fiberoptic Endoscopic Evaluation of Swallowing and the Swallowing Disturbances Questionnaire. CNS Spectr 24, 413–418 (2019).

33. Simpson, J.A., Lovecky, D., Kogan, J., Vetter, L.A. & Yohrling, G.J. Survey of the Huntington’s Disease Patient and Caregiver Community Reveals Most Impactful Symptoms and Treatment Needs. J Huntingtons Dis 5, 395–403 (2016).

34. Beglinger, L.J. et al. Earliest functional declines in Huntington disease. Psychiatry Res 178, 414–8 (2010).

35. Mason, S.L., Schaepers, M. & Barker, R.A. Problems with Social Cognition and Decision-Making in Huntington’s Disease: Why Is it Important? Brain Sci 11 (2021).

36. Barker, M.S. et al. Recognition memory and divergent cognitive profiles in prodromal genetic frontotemporal dementia. Cortex 139, 99–115 (2021).

37. Martinez-Horta, S.e.a. Neuropsychiatric symptoms are very common in premanifest and early stage Huntington’s Disease. Parkinsonism Relat Disord 25, 58–64 (2016).

38. van Duijn, E. et al. Neuropsychiatric symptoms in a European Huntington’s disease cohort (REGISTRY). J Neurol Neurosurg Psychiatry 85, 1411–8 (2014).

39. Andrews, S.C. et al. Apathy predicts rate of cognitive decline over 24 months in premanifest Huntington’s disease. Psychol Med, 1–7 (2020).

40. Migliore, S. et al. Cognitive and behavioral associated changes in manifest Huntington disease: A retrospective cross-sectional study. Brain Behav 11, e02151 (2021).

41. Galts, C.P.C. et al. Depression in neurodegenerative diseases: Common mechanisms and current treatment options. Neurosci Biobehav Rev 102, 56–84 (2019).

42. Kume, K. et al. Middle-age-onset cerebellar ataxia caused by a homozygous TWNK variant: a case report. BMC Med Genet 21, 68 (2020).

43. Henao, A.I. et al. Characteristic brain MRI findings in ataxia-neuropathy spectrum related to *POLG* mutation. Neuroradiol J 29, 46–8 (2016).

44. Kim, J.S. Delayed onset mixed involuntary movements after thalamic stroke: clinical, radiological and pathophysiological findings. Brain 124, 299–309 (2001).

45. Fukui, T. et al. Hemiballism-hemichorea induced by subcortical ischemia. Can J Neurol Sci 20, 324–8 (1993).

46. Den Dunnen, W.F.A. Trinucleotide repeat disorders. Handb Clin Neurol 145, 383–391 (2017).

47. Amlang, C.J., Trujillo Diaz, D. & Louis, E.D. Essential Tremor as a “Waste Basket” Diagnosis: Diagnosing Essential Tremor Remains a Challenge. Front Neurol 11, 172 (2020).

48. Anderson, D.G. et al. Comparison of the Huntington’s Disease like 2 and Huntington’s Disease Clinical Phenotypes. Mov Disord Clin Pract 6, 302–311 (2019).

49. Diehl, S.K. et al. Motor speech patterns in Huntington disease. Neurology 93, e2042–e2052 (2019).

50. Gagnon, M., Barrette, J. & Macoir, J. Language Disorders in Huntington Disease: A Systematic Literature Review. Cogn Behav Neurol 31, 179–192 (2018).

51. Bellosta Diago, E. et al. Circadian rhythm and autonomic dysfunction in presymptomatic and early Huntington’s disease. Parkinsonism Relat Disord 44, 95–100 (2017).

52. van Duijn, E. et al. Course of irritability, depression and apathy in Huntington’s disease in relation to motor symptoms during a two-year follow-up period. Neurodegener Dis 13, 9–16 (2014).

53. Julayanont, P., Heilman, K.M. & McFarland, N.R. Early-Motor Phenotype Relates to Neuropsychiatric and Cognitive Disorders in Huntington’s Disease. Mov Disord 35, 781–788 (2020).

54. Coleman, J.R.I. Shared Genetic Risk Between Psychiatric and Cognitive Symptoms in Huntington’s Disease and in the General Population. Biol Psychiatry 87, e25–e27 (2020).

55. Stevanin, G. et al. Huntington’s disease-like phenotype due to trinucleotide repeat expansions in the TBP and JPH3 genes. Brain 126, 1599–603 (2003).

56. Koutsis, G. et al. Genetic screening of Greek patients with Huntington’s disease phenocopies identifies an SCA8 expansion. J Neurol 259, 1874–8 (2012).

57. Sulek-Piatkowska, A. et al. Searching for mutation in the JPH3, ATN1 and TBP genes in Polish patients suspected of Huntington’s disease and without mutation in the IT15 gene. Neurol Neurochir Pol 42, 203–9 (2008).

58. Keckarevic, M. et al. Yugoslav HD phenocopies analyzed on the presence of mutations in PrP, ferritin, and Jp-3 genes. Int J Neurosci 115, 299–301 (2005).

59. Beck, J. et al. Large C9orf72 hexanucleotide repeat expansions are seen in multiple neurodegenerative syndromes and are more frequent than expected in the UK population. Am J Hum Genet 92, 345–53 (2013).

60. Beck, J. et al. Validation of next-generation sequencing technologies in genetic diagnosis of dementia. Neurobiol Aging 35, 261–5 (2014).

61. Renton, A.E. et al. A hexanucleotide repeat expansion in C9ORF72 is the cause of chromosome 9p21-linked ALS-FTD. Neuron 72, 257–68 (2011).

62. Ebbert, M.T.W. et al. Long-read sequencing across the C9orf72 ‘GGGGCC repeat expansion: implications for clinical use and genetic discovery efforts in human disease. Mol Neurodegener 13, 46 (2018).

63. Ibanez, K. et al. Whole genome sequencing for the diagnosis of neurological repeat expansion disorders in the UK: a retrospective diagnostic accuracy and prospective clinical validation study. Lancet Neurol 21, 234–245 (2022).

