## Supplemental Table 1 and Supplemental Table 2 for "Huntington’s disease phenocopy syndromes revisited: a clinical comparison and next-generation sequencing exploration"

### Supplementary Data

Table S-1: Survey questions sent to HD experts

| | | | |
|---|---|---|---|
| **Q1. What is your professional background? Please state your specialty.** | | | |
| Neurology | 36 / 70.6% | Movement Disorders | 25 / 49% |
| Neurogenetics | 11 / 21.6% | Psychiatry | 5 / 9.8% |
| Clinical Genetics | 1 / 2% | | |
| Answered | 51 | | |
| **Q2. How many years have you been working with HD and HD phenocopy patients?** | | | |
| 3 years | 1 / 1.9% | 17 years | 1 / 1.9% |
| 4 years | 1 / 1.9% | 19 years | 2 / 3.8% |
| 6 years | 2 / 3.8% | 20 years | 9 / 17.3% |
| 7 years | 1 / 1.9% | 22 years | 1 / 1.9% |
| 8 years | 2 / 3.8% | 23 years | 2 / 3.8% |
| 10 years | 4 / 7.7% | 25 years | 4 / 7.7% |
| 11 years | 3 / 5.8% | 28 years | 1 / 1.9% |
| 12 years | 1 / 1.9% | 30 years | 6 / 11.5% |
| 13 years | 1 / 1.9% | 31 years | 1 / 1.9% |
| 14 years | 2 / 3.8% | 34 years | 1 / 1.9% |
| 15 years | 5 / 9.6% | 36 years | 1 / 1.9% |
| Answered | 52 | | |
| **Q3. How many HD and/or HD phenocopy patients do you typically see in a month?** | | | |
| 1 patient | 3 / 5.8% | 18 patients | 2 / 3.8% |
| 2 patients | 1 / 1.9% | 20 patients | 5 / 9.6% |
| 4 patients | 2 / 3.8% | 25 patients | 4 / 7.7% |
| 5 patients | 2 / 3.8% | 30 patients | 2 / 3.8% |
| 6 patients | 2 / 3.8% | 35 patients | 1 / 1.9% |
| 8 patients | 3 / 5.8% | 40 patients | 4 / 7.7% |
| 10 patients | 12 / 23.1% | 45 patients | 1 / 1.9% |
| 12 patients | 1 / 1.9% | 50 patients | 1 / 1.9% |
| 15 patients | 4 / 7.7% | | |
| Answered | 52 | | |
| **Q4. In a patient with a compatible syndrome and a possibly positive family history (but without a genetic diagnosis in the family), what symptom or symptoms would make you most strongly consider an HD test?** | | | |
| Chorea | 52 / 100% | Agitation | 12 / 23.1% |
| Dystonia | 20 / 38.5% | Apathy | 28 / 53.9% |
| Rigidity | 9 / 17.3% | Anxiety | 17 / 32.7% |
| Gait abnormalities / Falls | 30 / 57.7% | Depression | 20 / 38.5% |
| Ataxia | 6 / 11.5% | Irritability | 34 / 65.4% |
| Tremor | 0 / 0% | Disinhibition | 16 / 30.8% |
| Dysarthria | 12 / 23.1% | Obsessive behaviour | 12 / 23.1% |
| Dysphagia / Choking | 11 / 21.2% | Paranoia | 5 / 9.6% |
| Memory loss | 9 / 17.3% | Delusions | 4 / 7.7% |
| Disorientation / Navigational difficulties | 5 / 9.6% | Hallucinations | 2 / 3.9% |

| | | | |
|---|---|---|---|
| Cognitive slowing | 25 / 48.1% | Change in dietary habits | 0 / 0% |
| Dysexecutive syndrome | 31 / 59.6% | Limb weakness | 0 / 0% |
| Loss of empathy | 13 / 25% | Weight loss | 11 / 21.2% |
| Insomnia | 4 / 7.7% | Pain | 0 / 0% |
| Hypersomnia | 0 / 0% | Neuropathy (sensory / motor) | 0 / 0% |
| Answered | 52 | | |
| **Q5. Based on Q4, please tick only your top 3 symptom which would make you consider an HD test.** | | | |
| Chorea | 52 / 100% | Agitation | 0 / 0% |
| Dystonia | 13 / 25% | Apathy | 6 / 11.5% |
| Rigidity | 1 / 1.9% | Anxiety | 2 / 3.9% |
| Gait abnormalities / Falls | 15 / 28.9% | Depression | 2 / 3.9% |
| Ataxia | 2 / 3.9% | Irritability | 16 / 30.8% |
| Tremor | 0 / 0% | Disinhibition | 6 / 11.5% |
| Dysarthria | 1 / 1.9% | Obsessive behaviour | 0 / 0% |
| Dysphagia / Choking | 2 / 3.9% | Paranoia | 1 / 1.9% |
| Memory loss | 3 / 5.8% | Delusions | 0 / 0% |
| Disorientation / Navigational difficulties | 0 / 0% | Hallucinations | 0 / 0% |
| Cognitive slowing | 16 / 30.8% | Change in dietary habits | 0 / 0% |
| Dysexecutive syndrome | 16 / 30.8% | Limb weakness | 0 / 0% |
| Loss of empathy | 1 / 1.9% | Weight loss | 1 / 1.9% |
| Insomnia | 1 / 1.9% | Pain | 0 / 0% |
| Hypersomnia | 0 / 0% | Neuropathy (sensory / motor) | 0 / 0% |
| Answered | 52 | | |
| **Q6. In such a patient, is there a clinical symptom (or symptoms) that, if present, would make you less inclined to order an HD test, possibly because that symptom is making another diagnosis more likely in your mind?** | | | |
| Chorea | 1 / 2% | Agitation | 1 / 2% |
| Dystonia | 1 / 2% | Apathy | 0 / 0% |
| Rigidity | 5 / 9.8% | Anxiety | 2 / 3.9% |
| Gait abnormalities / Falls | 2 / 3.9% | Depression | 3 / 5.9% |
| Ataxia | 22 / 43.1% | Irritability | 0 / 0% |
| Tremor | 25 / 49% | Disinhibition | 4 / 7.8% |
| Dysarthria | 2 / 3.9% | Obsessive behaviour | 1 / 2% |
| Dysphagia / Choking | 1 / 2% | Paranoia | 0 / 0% |
| Memory loss | 7 / 13.7% | Delusions | 0 / 0% |
| Disorientation / Navigational difficulties | 5 / 9.8% | Hallucinations | 13 / 25.5% |
| Cognitive slowing | 1 / 2% | Change in dietary habits | 6 / 11.8% |
| Dysexecutive syndrome | 0 / 0% | Limb weakness | 30 / 58.8% |
| Loss of empathy | 1 / 2% | Weight loss | 0 / 0% |
| Insomnia | 3 / 5.9% | Pain | 28 / 54.9% |
| Hypersomnia | 7 / 13.7% | Neuropathy (sensory / motor) | 38 / 74.5% |
| Answered | 51 | | |
| **Q7. Based on Q6, please tick only your 3 top symptom choices that would make you less inclined to order an HD test.** | | | |
| Chorea | 0 / 0% | Agitation | 0 / 0% |
| Dystonia | 0 / 0% | Apathy | 0 / 0% |
| Rigidity | 1 / 2% | Anxiety | 1 / 2% |

| | | | |
|---|---|---|---|
| Gait abnormalities / Falls | 0 / 0% | Depression | 1 / 2% |
| Ataxia | 18 / 35.3% | Irritability | 0 / 0% |
| Tremor | 14 / 27.5% | Disinhibition | 1 / 2% |
| Dysarthria | 0 / 0% | Obsessive behaviour | 0 / 0% |
| Dysphagia / Choking | 1 / 2% | Paranoia | 0 / 0% |
| Memory loss | 4 / 7.8% | Delusions | 0 / 0% |
| Disorientation / Navigational difficulties | 3 / 5.9% | Hallucinations | 11 / 21.6% |
| Cognitive slowing | 0 / 0% | Change in dietary habits | 6 / 11.8% |
| Dysexecutive syndrome | 0 / 0% | Limb weakness | 28 / 54.9% |
| Loss of empathy | 0 / 0% | Weight loss | 1 / 2% |
| Insomnia | 1 / 2% | Pain | 17 / 33.3% |
| Hypersomnia | 3 / 5.9% | Neuropathy (sensory / motor) | 38 / 74.5% |
| Answered | 51 | | |
| **Q8. What combination of symptoms would be sufficient for you to consider HD and test for it?** | | | |
| Chorea - cognitive problems - anxiety - with family history (one affected parent or grandparent) | | | 48 / 92.3% |
| Chorea - cognitive problems - anxiety - no family history | | | 43 / 82.7% |
| Dystonia - cognitive problems - anxiety - with family history (one affected parent or grandparent) | | | 36 / 69.2% |
| Dystonia - cognitive problems - anxiety - no family history | | | 19 / 36.5% |
| Rigidity- cognitive problems - apathy- with family history (one affected parent or grandparent) | | | 31 / 59.6% |
| Rigidity- cognitive problems - apathy- no family history | | | 11 / 21.2% |
| Chorea - with family history (one affected parent or grandparent) | | | 46 / 88.5% |
| Chorea - no family history | | | 36 / 69.2% |
| Dystonia - with family history (one affected parent or grandparent) | | | 27 / 51.9% |
| Dystonia - no family history | | | 3 / 5.8% |
| Rigidity - with family history (one affected parent or grandparent) | | | 25 / 48.1% |
| Rigidity - no family history | | | 2 / 3.9% |
| Dysexecutive syndrome - with family history (one affected parent or grandparent) | | | 29 / 55.8% |
| Dysexecutive syndrome - no family history | | | 2 / 3.9% |
| Irritability - cognitive slowing - with family history (one affected parent or grandparent) | | | 34 / 65.4% |
| Irritability - cognitive slowing - no family history | | | 8 / 15.4% |
| Dysexecutive syndrome - depression - with family history (one affected parent or grandparent) | | | 30 / 57.7% |
| Dysexecutive syndrome - depression - no family history | | | 2 / 3.9% |
| Disinhibition - with family history (one affected parent or grandparent) | | | 20 / 38.5% |
| Disinhibition - no family history | | | 1 / 1.9% |
| Answered | | | 52 |
| **Q9. Are there any symptoms that, if present, would make you expect that an HD test will come back negative, despite the obvious need to test for (and exclude) HD?** | | | |
| Chorea | 0 / 0% | Agitation | 0 / 0% |
| Dystonia | 1 / 1.9% | Apathy | 0 / 0% |
| Rigidity | 0 / 0% | Anxiety | 1 / 1.9% |
| Gait abnormalities / Falls | 0 / 0% | Depression | 1 / 1.9% |
| Ataxia | 11 / 21.2% | Irritability | 0 / 0% |
| Tremor | 13 / 25% | Disinhibition | 0 / 0% |
| Dysarthria | 1 / 1.9% | Obsessive behaviour | 0 / 0% |

| | | | |
|---|---|---|---|
| Dysphagia / Choking | 0 / 0% | Paranoia | 1 / 1.9% |
| Memory loss | 4 / 7.7% | Delusions | 1 / 1.9% |
| Disorientation / Navigational difficulties | 2 / 3.9% | Hallucinations | 6 / 11.5% |
| Cognitive slowing | 0 / 0% | Change in dietary habits | 3 / 5.8% |
| Dysexecutive syndrome | 1 / 1.9% | Limb weakness | 24 / 46.2% |
| Loss of empathy | 0 / 0% | Weight loss | 1 / 1.9% |
| Insomnia | 1 / 1.9% | Pain | 21 / 40.4% |
| Hypersomnia | 9 / 17.3% | Neuropathy (sensory / motor) | 32 / 61.5% |
| None of these | 12 / 23.1% | | |
| Answered | 52 | | |
| **Q10. Based on Q9, which top 3 symptoms would make you expect that an HD test will come back negative, despite the obvious need to test for it?** | | | |
| Chorea | 0 / 0% | Agitation | 0 / 0% |
| Dystonia | 1 / 1.9% | Apathy | 0 / 0% |
| Rigidity | 0 / 0% | Anxiety | 1 / 1.9% |
| Gait abnormalities / Falls | 0 / 0% | Depression | 1 / 1.9% |
| Ataxia | 8 / 15.4% | Irritability | 0 / 0% |
| Tremor | 10 / 19.2% | Disinhibition | 0 / 0% |
| Dysarthria | 0 / 0% | Obsessive behaviour | 0 / 0% |
| Dysphagia / Choking | 0 / 0% | Paranoia | 1 / 1.9% |
| Memory loss | 2 / 3.9% | Delusions | 1 / 1.9% |
| Disorientation / Navigational difficulties | 2 / 3.9% | Hallucinations | 4 / 7.7% |
| Cognitive slowing | 0 / 0% | Change in dietary habits | 3 / 5.8% |
| Dysexecutive syndrome | 1 / 1.9% | Limb weakness | 24 / 46.2% |
| Loss of empathy | 0 / 0% | Weight loss | 2 / 3.9% |
| Insomnia | 0 / 0% | Pain | 18 / 34.6% |
| Hypersomnia | 2 / 3.9% | Neuropathy (sensory / motor) | 34 / 65.4% |
| None of these | 10 / 19.2% | | |
| Answered | 52 | | |

##### Table S-2: Details of patient samples selected for whole-genome sequencing

Based on the selection criteria described in Table 1, 50 samples were selected for whole-genome sequencing; their details are described in this table. Sex, age at onset (AAO) and the strength of the family history (Goldman score, see Figure II-1) were noted, and the clinical notes scoured for symptoms the patients developed in their lifetime, as well as the last time they presented in clinic at the National Hospital for Neurology and Neurosurgery (NHNN). Based on this information and their respective score, patient samples were shortlisted and selected. HDPS Movement + HDPS Movement Disorder; HDPS Cognitive = HDPS Cognitive Decline; HDPS Psychiatric = HDPS Psychiatric Disturbance; Neuropath = Neuropathology available

| HDPC Sample | Sex | AAO | Goldman Score | HDPS Movement | HDPS Cognitive | HDPS Psychiatric | HDPS Total | Neuro-path | Chorea present | Last Clinic (years) | Total score |
|---|---|---|---|---|---|---|---|---|---|---|---|
| **1** | Fe | 43 | 1 | 1 | 1 | 1 | 3 | no | yes | 2 | 85 |
| **2** | Fe | 55 | 1 | 1 | 1 | 1 | 3 | no | yes | 2 | 85 |
| **3** | Mal | 59 | 1 | 1 | 1 | 1 | 3 | no | no | 5 | 81 |
| **4** | Mal | 61 | 1 | 1 | 1 | 0 | 2 | yes | no | 2 | 80 |
| **5** | Fe | 69 | 2 | 1 | 1 | 1 | 3 | no | yes | 2 | 75 |
| **6** | Fe | 45 | 2 | 1 | 1 | 1 | 3 | no | yes | 2 | 75 |
| **7** | Mal | 50 | 3.5 | 1 | 1 | 1 | 3 | no | yes | 2 | 67 |
| **8** | Fe | 49 | 3.5 | 1 | 1 | 1 | 3 | no | yes | 2 | 67 |
| **9** | Mal | 59 | 4 | 1 | 1 | 1 | 3 | no | yes | 2 | 65 |
| **10** | Mal | 53 | 4 | 1 | 1 | 1 | 3 | no | no | 2 | 65 |
| **11** | Mal | 70 | 4 | 1 | 1 | 1 | 3 | no | yes | 2 | 65 |
| **12** | Fe | 39 | 1 | 1 | 1 | 0 | 2 | no | yes | 2 | 65 |
| **13** | Fe | 49 | 4.5 | 1 | 1 | 1 | 3 | no | no | 2 | 65 |
| **14** | Mal | 43 | 4 | 1 | 1 | 1 | 3 | no | yes | 2 | 65 |
| **15** | Mal | 64 | 4 | 1 | 1 | 1 | 3 | no | yes | 2 | 65 |
| **16** | Fe | 11 | 1 | 1 | 0 | 1 | 2 | no | yes | 2 | 65 |
| **17** | Mal | 43 | 3.5 | 1 | 1 | 1 | 3 | no | no | 3 | 64 |
| **18** | Mal | 53 | 3.5 | 1 | 1 | 1 | 3 | no | no | 53 | 63 |
| **19** | Mal | 39 | 3.5 | 1 | 1 | 1 | 3 | no | no | - | 62 |
| **20** | Mal | 47 | 4 | 1 | 1 | 1 | 3 | no | no | 3 | 62 |
| **21** | Fe | 43 | 1 | 0 | 1 | 1 | 2 | no | no | - | 60 |
| **22** | Fe | 69 | 4 | 1 | 1 | 1 | 3 | no | yes | - | 60 |
| **23** | Mal | 33 | 4 | 1 | 1 | 1 | 3 | no | yes | - | 60 |
| **24** | Fe | 34 | 2 | 1 | 0 | 1 | 2 | no | no | 2 | 55 |
| **25** | Fe | 55 | 2 | 1 | 1 | 0 | 2 | no | no | 2 | 55 |
| **26** | Mal | 75 | 3.5 | 1 | 1 | 0 | 2 | no | yes | 2 | 47 |
| **27** | Fe | 63 | 3.5 | 1 | 1 | 0 | 2 | no | yes | 2 | 47 |
| **28** | Fe | 66 | 3.5 | 1 | 0 | 1 | 2 | no | yes | 2 | 47 |

| HDPC Sample | Sex | AAO | Goldman Score | HDPS Movement | HDPS Cognitive | HDPS Psychiatric | HDPS Total | Neuro-path | Chorea present | Last Clinic (years) | Total score |
|---|---|---|---|---|---|---|---|---|---|---|---|
| **29** | Mal | 44 | 3.5 | 1 | 1 | 0 | 2 | no | no | 2 | 47 |
| **30** | Fe | 62 | 3.5 | 1 | 1 | 0 | 2 | no | no | 2 | 47 |
| **31** | Fe | 34 | 3.5 | 1 | 0 | 1 | 2 | no | yes | 2 | 47 |
| **32** | Fe | 70 | 3.5 | 1 | 0 | 1 | 2 | no | yes | 2 | 47 |
| **33** | Fe | 65 | 1 | 1 | 1 | 0 | 2 | no | no | 5 | 46 |
| **34** | Mal | 16 | 4 | 0 | 1 | 1 | 2 | no | no | 2 | 45 |
| **35** | Fe | 68 | 4 | 1 | 0 | 1 | 2 | no | yes | 2 | 45 |
| **36** | Fe | 46 | 3 | 1 | 1 | 0 | 2 | no | no | 5 | 45 |
| **37** | Mal | 77 | 4 | 1 | 1 | 0 | 2 | no | yes | 2 | 45 |
| **38** | Mal | 51 | 4 | 1 | 1 | 0 | 2 | no | yes | 2 | 45 |
| **39** | Mal | 64 | 4 | 1 | 1 | 0 | 2 | no | yes | 2 | 45 |
| **40** | Fe | 53 | 4 | 1 | 1 | 0 | 2 | no | yes | 2 | 45 |
| **41** | Fe | 46 | 4 | 1 | 1 | 0 | 2 | no | no | 2 | 45 |
| **42** | Fe | 53 | 4 | 1 | 1 | 0 | 2 | no | yes | 2 | 45 |
| **43** | Fe | 58 | 4 | 1 | 1 | 0 | 2 | no | no | 2 | 45 |
| **44** | Fe | 22 | 1 | 1 | 0 | 0 | 1 | no | yes | 2 | 35 |
| **45** | Fe | 52 | 1 | 1 | 0 | 0 | 1 | no | yes | 2 | 35 |
| **46** | Fe | 52 | 1 | 1 | 0 | 0 | 1 | no | yes | 2 | 35 |
| **47** | Mal | 40 | 1 | 1 | 0 | 0 | 1 | no | no | 2 | 35 |
| **48** | Mal | 67 | 1 | 0 | 1 | 0 | 1 | no | no | 2 | 35 |
| **49** | Fe | 35 | 1 | 1 | 0 | 0 | 1 | no | no | - | 30 |
| **50** | Fe | 45 | 1 | 0 | 0 | 0 | 0 | 0 | no | 5 | 21 |
